# Modeling the contribution of antibodies to the within-host dynamics of single and dual helminth infections in a natural system

**DOI:** 10.1101/2022.10.20.513085

**Authors:** Chiara Vanalli, Lorenzo Mari, Renato Casagrandi, Brian Boag, Marino Gatto, Isabella M. Cattadori

## Abstract

Within-host models of infection can provide important insights into the processes that affect parasite spread and persistence in host populations. However, modeling is often limited by the availability of empirical data, a problem commonly encountered in natural systems. Here, we used six years of immune-infection observations of two gastrointestinal helminths (*Trichostrongylus retortaeformis* and *Graphidium strigosum*) from a population of European rabbits (*Oryctolagus cuniculus*) to develop an age-dependent, within-host mathematical model that explicitly included species-specific and cross-reacting antibody (IgA and IgG) responses to each helminth in hosts with single or dual infections. Different models of single infection were formally compared to test alternative mechanisms of parasite regulation. The two models that best described single infections of each helminth species were then coupled through antibody cross-immunity to examine how the presence of one species could alter the host immune response to, and the within-host dynamics of, the other species. For both single infections, model selection suggested that either IgA or IgG responses could equally explain the observed parasite intensities at different host age. However, the strength of this response drastically changed between the two helminths, being stronger against *T. retortaeformis* than against *G. strigosum* and causing contrasting age-intensity profiles. When the two helminths coinfect the same host, we found a decrease of the species-specific immune response to both species together with an asymmetric cross-immune response driven by IgG. Changes in the level and affinity of antibodies from single to dual infections contributed to the significant increase of both helminth abundances. By combining mathematical modeling with immuno-infection data, our work provides a model framework for disentangling some of the complexities generated by host-parasite and parasite-parasite interactions from natural systems. Our approach thus offers a tractable general tool to examine immune-infection relationships within hosts.

**Author summary:** The host immune response often plays a critical role in regulating parasite dynamics and transmission. We developed a mathematical model to evaluate whether and how the observed variation in host antibody responses could explain the relative differences in the abundance of two parasite species in hosts with single and dual infections from a natural rabbit population. Our results indicated that either IgA or IgG could describe the contrasting trends of the two parasites in rabbits with single infections. Specifically, antibodies appeared to control *T. retortaeformis*, while the effect was less clear for *G. strigosum*. For dual infections, we found a weaker specific antibody response against both helminths and an asymmetric cross-immunity, which could explain the significantly greater intensities observed for *T. retortaeformis* and, secondly, for *G. strigosum*. Our within-host mathematical framework provides a plausible mechanism for the mediated role of antibodies in hosts with single and dual parasite infections, and how the strength of these infection-immune interactions changes with host age. This model framework offers a way forward to our understanding of the within-host processes that generate individual variation in infection and is relatively flexible to be applied to other natural systems.

## Introduction

The study of the dynamics of infectious diseases in natural systems has been primarily applied at the level of host populations, and mathematical models have been very successful at capturing the fundamental processes of infection and onward transmission, including where and when heterogeneities in these processes typically emerge (1–5). The within-host modeling of infections have received less attention, although their popularity has rapidly increased, mostly facilitated by the advancement of molecular tools that have allowed the quantification of the infection-immune interactions at the cellular and molecular level (6–9). These models primarily centered around microparasitic infections of humans and the description of their control by the host immune response, vaccines or drug treatments, notably for HIV (10,11), dengue (12), influenza (13), malaria (14), and more recently SARS-CoV-2 (15).

The within-host modeling of macroparasitic infections, primarily helminths, has been largely overlooked, although several classical theoretical (16,17) and phenomenological studies (18–22) have examined within-host processes as nested components of epidemiological models of infection and transmission. These model frameworks have conceptually incorporated acquired immunity as a function of host exposure to past and current infections by the target parasite, quantified as accumulated parasite burden through the host life span (20,22,23). For example, this approach was used to describe the development and wane of the host immune response against trichostrongylid infections in domestic ruminants where animals were exposed to free-living infective stages and subsequently lost immune memory (24,25). Austin and Anderson (1996) (26) further developed this concept by describing the dynamics of acquired immunity to an infection as a predator-prey mechanistic interaction between host’s antibodies and parasite’s antigens. The general models that were originally proposed have been further developed and tailored to specific systems and goals, for example, they have been used to evaluate the effect of immunity on the establishment of infective larvae or on the survival and fecundity of adult parasites (16,18,20,27), including the protective effect of immunity to re-infections (22,28).

While these approaches have provided a tractable framework for the interactions between host immunity and helminth infections, the quantitative understanding of these relationships using empirical data and mathematical modeling is still limited. What makes these within-host modeling notably challenging is the often-complex life cycle of the parasite and the associated host immune response (29,30). This is even more problematic for natural systems that require additional efforts and tools for data collection and sample processing, and where system heterogeneities contribute to complexities in host-parasite interactions that can be difficult to disentangle.

A frequent source of host heterogeneity in natural systems is the occurrence of co-infections, individuals that carry more than one parasite species, or parasite strain, at any one time in their life (31,32). The presence of a second parasite species can alter the host immune response to the target species (33–36) and facilitate or constrain its dynamics of infection and life history (37–39). These interspecific interactions have been modeled by borrowing again the predator-prey concept, where a single generalist predator, represented by host immunity, can target several preys represented by the community of coinfecting parasites (40–42). By mediating the interactions between parasite species, the immune response can contribute to processes of competition (36,43) or facilitation (44–46) within individual hosts, and in turn affect the transmission of the coinfecting species at the host population level. The study of the immune-mediated mechanisms of interaction between helminths has strengthened our understanding of the dynamics of coinfection (47), however, the design and implementation of models contrasted to empirical data from field systems has been largely neglected.

We used observations from a natural population of European rabbit (*Oryctolagus cuniculus*) infected with one or both the gastrointestinal helminths *Trichostrongylus retortaeformis* and *Graphidium strigosum* to develop a within-host mathematical model aimed at examining how antibodies (IgA and IgG) regulated the intensity of infection for each helminth and at assessing how these mechanisms changed in hosts with both helminths. Different hypotheses were tested to investigate alternative mechanisms of antibody regulation, including the mediated role in the interactions between the two helminths. Our aim was to develop a within-host model that could track changes of antibody responses and intensities of infection from field observations, and to compare the findings with previous knowledge on this host-parasite system from laboratory experiments (48–50).

## Materials and Methods

### The helminth-rabbit system

*Trichostrongylus retortaeformis* and *Graphidium strigosum* are common gastrointestinal helminths that cause chronic infections in the European rabbit (*Oryctolagus cuniculus*) (51–53) by ingestion of free-living third-stage larvae (L3) from the pasture. *G. strigosum* settles in the stomach while *T. retortaeformis* in the small intestine, where they then develop into adults, reproduce and shed eggs back onto the pasture through the rabbit’s feces (49,50). The two helminths are often found coinfecting the same rabbit (51,54) with intensities that vary differently with host age and seasonal trends (46,52,55).

Previous studies from natural rabbit populations (51,52), including data used for the current work (54), showed that *T. retortaeformis* intensities peak in young rabbits and decrease in older hosts, while *G. strigosum* intensities accumulate with host age with weak or no evidence of parasite control. These different dynamics have been suggested to be regulated by host immunity, which appears to be more effective against the former than the latter helminth, although does not protect from reinfection (51,52,55,56). Laboratory experiments confirmed the important role of host immunity. For example, rabbits developed an anti-inflammatory Th-2 immune response against both helminths, which involved the production of IL4 and IL13 cytokines, IgA and IgG antibodies, and eosinophils (48–50,57). Specifically, the two antibodies showed contrasting trends with host age, and a stronger response was found against *T. retortaeformis* than *G. strigosum* (50). This result was supported by experimental reinfections after anthelminthic treatment, where the IgA immune reaction was more apparent against *T. retortaeformis* than *G. strigosum* (57). More generally, antibodies contribute to the humoral response to gut infections (58,59) and often act as one of the main regulatory factors of parasite intensity by affecting mortality or expulsion (60–62), and fecundity or egg shedding (63,64). The down-modulation of the immune response by *G. strigosum* was also suggested as a possible mechanism that facilitated the accumulation of this helminth and, in turn, could have also contributed to the higher intensities of *T. retortaeformis* in co-infected rabbits, when compared to hosts with single infections (32). More recently, the modeling of these two helminths from this dataset (including the population of the current study) supported the role of IgA-mediated facilitation as a plausible mechanism that could explain the higher intensities of the two helminths in co-infected rabbits (46). A more complex model framework centered around the IL4-IgA immune interaction and data from laboratory experiments showed that, although the two helminths stimulated a similar immune response, changes in IgA and IL4 levels could describe the rapid clearance observed for *T. retortaeformis* but not *G. strigosum* when compared to single infections (48,50).

To explore in more detail the role of antibodies in the within-host dynamics of *T. retortaeformis* and *G. strigosum*, we further elaborated on previous modelling frameworks by focusing on a natural population of rabbits from Perthshire (Scotland, UK), for which individual data on the intensity of infection and associated specific IgA and IgG responses were available monthly from 2005 to 2010. For every rabbit, the intensity of the two helminths was quantified by aliquots following standardized parasitological techniques (54). Specific IgA and IgG responses against excretory/secretory products of adult parasites of each species were available from blood serum collected at the time of sampling (54), quantified via the Enzyme-Linked Immunosorbent Assay (ELISA) and presented as Optical Density Index (ODI) (50). Rabbit age was determined by classifying the animals in eight monthly classes, from one to eight+ months old individual, based on total body mass and further confirmed using eye-lens mass (52,65).

For our analysis, rabbits were divided into three groups according to their infection status at the time of sampling: i) rabbits infected only with *T. retortaeformis* (TR, n=476), ii) rabbits infected only with *G. strigosum* (GS, n=254), and iii) rabbits infected with both helminths (TR-GS, n=910). Uninfected rabbits, which mainly belong to the younger age classes, were added to each infection group to represent baseline values (46). Rabbits that were found positive to myxomatosis were excluded from the dataset to avoid any interference caused by the virus infection (66). For practical modeling reasons, we assumed that these three groups do not affect each other dynamics. Our general aim was to evaluate the age-dependent, within-host processes of infection and antibody response and, thus, we ignored the possible interannual variations and grouped data collected in different years from hosts of the same age. Similarly, the consequences of these within-host interactions for helminth transmission at the host population level were not addressed as they were investigated elsewhere (46).

### Single infection: Model description, selection and calibration

For each helminth, we developed a within-host model of infection that explicitly tracks changes of parasite intensity (i.e. the individual parasite load), *P*_*i*_ (with *i*=TR for *T. retortaeformis* or GS for *G. strigosum*), and species-specific IgA and IgG levels, named as *A*_*i*_ and *G*_*i*_ respectively, at host age *a* as follow:

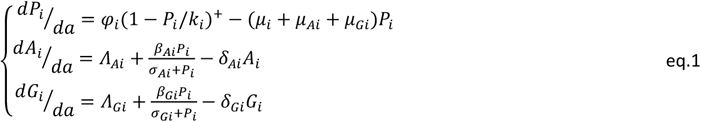

Rabbits become infected by ingesting third stage larvae (L3) of helminth *i* while foraging at an age-dependent rate *φ*_*i*_(*a*) = *ϕ*_*i*_((200+ 275a)/3340)^*γi*^ (55,56). The parameters *ϕ*_*i*_ and *γ*_*i*_ represent the baseline value of the feeding rate and its positive dependence upon host age, respectively, and together describe the rate at which rabbits are exposed to L3 larvae while feeding (67). We assumed that ingested larvae immediately develop into adult parasites and settle in their specific sites of infection according to a density-dependence regulation function, in which *k*_*i*_ represents the carrying capacity. Adult parasites die at a natural baseline rate *μ*_*i*_, fixed according to an average lifespan of the host of one year (68). Parasites can die at a faster rate because of antibody response, where *μ*_*Ai*_ and *μ*_*Gi*_ represent IgA- and IgG-induced parasite mortalities, respectively. We define these mortalities as the attack rates of antibodies to the parasites. Antibody attack rates are a function of the antibody level at a specific host age class, *A*_*i*_(*a*) (IgA O.D.Index) and *G*_*i*_(*a*) (IgG O.D.Index), and antibody affinity here quantified as their respective abilities to kill parasites by quickly binding to their antigens. For each helminth *i*, two different forms of antibody affinity were examined:

i. a constant affinity, where the parasite killing rate per unit of antibody is constant throughout the host life span, *α*_*Ai*_ and *α*_*Gi*_, with corresponding Ig attack rates that are proportional to antibody levels *μ*_*Ai*_ = *α*_*Ai*_ *A*_*i*_ and *μ*_*Gi*_ = *α*_*Gi*_*G*_*i*_;
ii. an increasing affinity, which is proportional to the accumulated host exposure to infective stages, namely, the integral of exposure to species *i* from the birth of the rabbit to its age *a* at the sampling, explicitly: 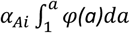 and 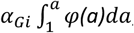, with resulting antibody attack rates 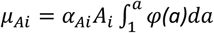 and 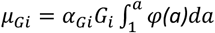 (20,22,23).

We note that, irrespective of our modelling assumption, there is no infection in the first month of age as newborn rabbits start moving from a milk to a grass diet towards the end of their first month of life. In absence of helminths (*P*_*i*_ =0), antibodies are maintained at equilibrium levels, *Ā*_*i*_ = *Λ*_*Ai*_/*δ*_*Ai*_ and 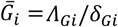 *Λ*_*Gi*_/*δ*_*Gi*_, which are calculated as ratios between the baseline production rates *Λ*_*Ai*_ and *Λ*_*Gi*_ and the decay rates *δ*_*Ai*_ and *δ*_*Gi*_, following Vanalli et al. (2020) (48). The IgA and IgG equilibrium levels were available from laboratory naїve rabbits (57). Laboratory experiments also provided the antibody decay rates *δ*_*Ai*_ and *δ*_*Gi*_ (6). We assumed that in presence of infection (*P*_*i*_ >0), *A*_*i*_ and *G*_*i*_ production is stimulated by parasite intensity according to a saturating function described by the coefficients *β*_*Ai*_, *σ*_*Ai*_ and *β*_*Gi*_, *σ*_*Gi*_ for IgA and IgG, respectively (49,50). The definition of the model parameters together with units of measures and credible values (whenever available) are reported in Table 1.

**Table 1.** Single infection: Model parameters, definitions, dimensions and available values. (Tbc=to be calibrated).

| Parameter | Definition | Unit of measure | *T. retortaeformis* | *G. strigosum* |
| --- | --- | --- | --- | --- |
| $\phi_i$ | Baseline feeding rate | parasites month<sup>-1</sup> rabbit<sup>-1</sup> | Tbc | Tbc |
| $\gamma_i$ | Allometric exponent of the host feeding rate | - | Tbc | Tbc |
| $k_i$ | Parasite carrying capacity | Parasites | Tbc | Tbc |
| $\mu_i$ | Natural parasite mortality rate | month<sup>-1</sup> | 8.22×10<sup>-2</sup> | |
| $\alpha_{Ai}$ | Parasite killing rate per unit of antibody | (ODI<sup>-1</sup> month<sup>-1</sup>) or (ODI<sup>-1</sup> month<sup>-1</sup> parasites<sup>-1</sup>) | Tbc | Tbc |
| $\alpha_{Gi}$ | Parasite killing rate per unit of IgG antibody | (ODI<sup>-1</sup> month<sup>-1</sup>) or (ODI<sup>-1</sup> month<sup>-1</sup> parasites<sup>-1</sup>) | Tbc | Tbc |
| $\bar{A}_i$ | IgA equilibrium value | ODI | 0.180 | 0.104 |
| $\Lambda_{Ai}$ | IgA baseline production rate | ODI month<sup>-1</sup> | 0.116 | 0.168 |
| $\delta_{Ai}$ | IgA natural decay rate | month<sup>-1</sup> | 0.646 | 1.62 |
| $\beta_{Ai}$ | IgA saturation value | ODI month<sup>-1</sup> | Tbc | Tbc |
| $\sigma_{Ai}$ | Parasite intensity at which IgA response is ½ of $\beta_{Ai}$ | Parasites | Tbc | Tbc |
| $\bar{G}_i$ | IgG equilibrium value | ODI | 3.23×10<sup>-2</sup> | 4.26×10<sup>-2</sup> |
| $\Lambda_{Gi}$ | IgG baseline production rate | ODI month<sup>-1</sup> | 5.94×10<sup>-2</sup> | 3.50×10<sup>-2</sup> |
| $\delta_{Gi}$ | IgG natural decay rate | month<sup>-1</sup> | 1.84 | 0.823 |
| $\beta_{Gi}$ | IgG saturation value | ODI month<sup>-1</sup> | Tbc | Tbc |
| $\sigma_{Gi}$ | Parasite intensity at which IgG response is ½ of $\beta_{Gi}$ | Parasites | Tbc | Tbc |

We tested different hypothesis-driven models of parasite regulation by examining the contribution of IgA, IgG, and parasite density-dependence to the intensity of infection, and how these interactions changed with host age. Three main hypotheses, and their related combinations, were investigated:

1. IgA regulates the intensity of parasite *i* via an attack rate *μ*_*Ai*_ ;
2. IgG regulates the intensity of parasite *i* via an attack rate *μ*_*Gi*_;
3. Density-dependence regulates the intensity of parasite *i* according to the parasite carrying capacity *k*_*i*_.

These three hypotheses generated 14 competing models that selectively evaluate different combinations of parasite regulatory mechanisms (Table 2). For sake of simplicity, mixed hypothesis on the two forms of attack rates by IgA, *μ*_*Ai*_, and IgG, *μ*_*Gi*_, were not examined and we assumed that they were either constant or the integral of host exposure, for both helminths.

**Table 2.**
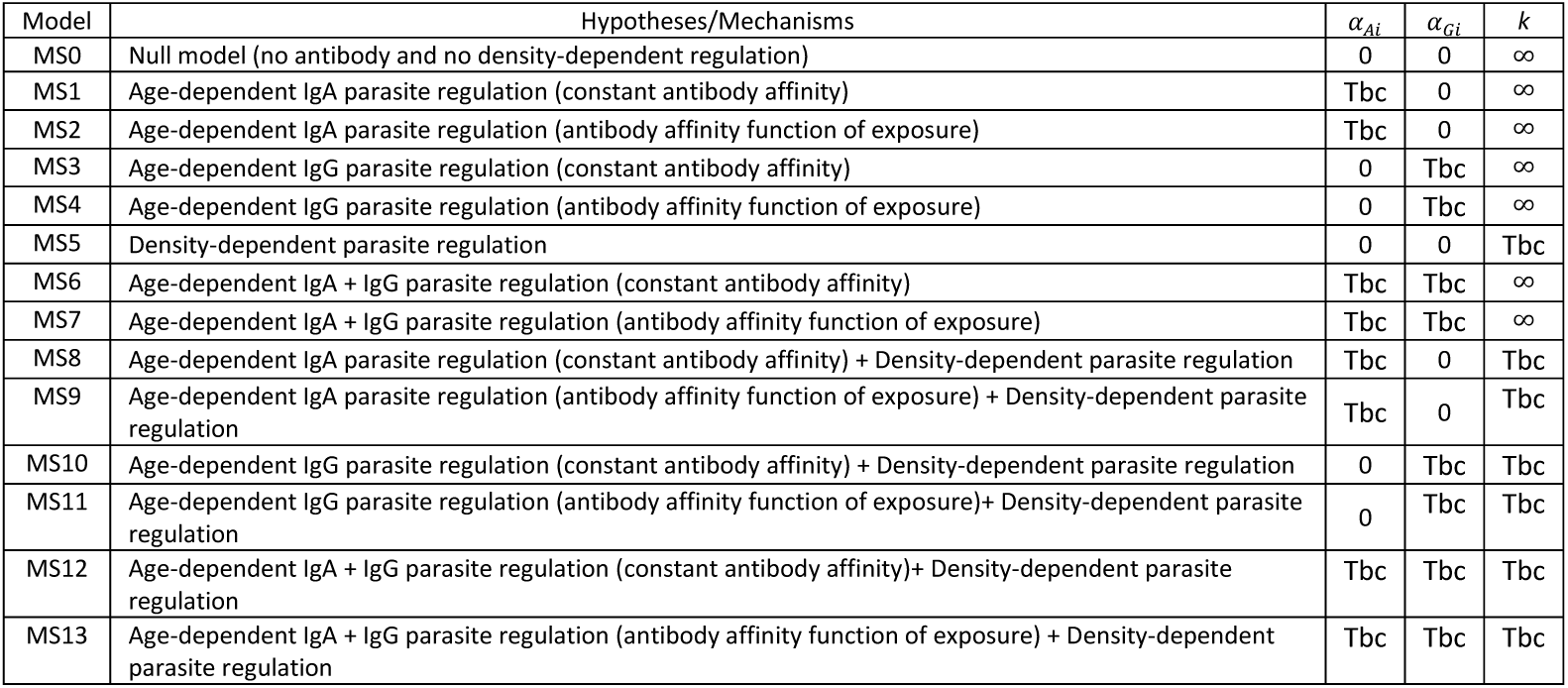
Tested hypotheses and related mechanisms for the competing models of single infection. The parameters *α*_*Ai*_, *α*_*Gi*_ and *k* are set equal to 0/∞ when the respective mechanism is not considered, otherwise they are calibrated (Tbc=to be calibrated).

Model calibration was performed, on every rabbit and independently for the three infection groups, by assuming that data are affected by lognormal noise and minimizing the loglikelihood, *ERR*, calculated as a weighted sum of logarithms of the logarithms of the variable errors, *ERR*_*P*_ (for parasite intensity), *ERR*_*A*_ (for IgA), and *ERR*_*G*_ (for IgG), see Appendix S1. Also, we considered the animal sample size, *n*_*j*_, of each of the *j*=1…8 monthly age classes, and calculated *ERR* as follows:

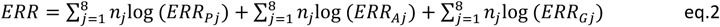

Since the variables are affected by lognormal noise and characterized by different magnitudes and the age classes have different sample size *n*_*j*_, we considered the percentage errors when comparing their errors (48). Each error component of age class *j* (eq. 2) was thus computed as the logarithmic square ratio between observed and estimated values, normalized by the data sample size:

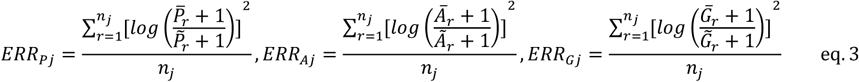

Here, 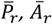 and 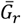 represent the observed parasite intensity and related IgA and IgG levels, respectively, for each rabbit *r* belonging to age class *j*, while 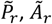 and 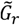 are the estimated values of the variables of interest. Model calibration was performed using MATLAB (R2019b), through the combination of the genetic algorithm from the Optimization Toolbox (*ga*) and a non-linear solver (*fminsearch*) to exclude local solutions during the optimization of the error function and thus to obtain a precise estimate of the parameters. We selected the best model, among the candidate set, independently for each helminth species and infection group, based on the best compromise between goodness of fit and parsimony, according to the Akaike Information Criterion (*AIC*). Specifically, for each model we evaluated the score *AIC* = *ERR* + 2*h*, where *ERR* represents the minimized loglikelihood (see eq. 2) and *h* is the model complexity, i.e. the number of parameters to calibrate (69). The model with the lowest AIC was selected.

### Dual infection: Model description, selection and calibration

For rabbits infected with both helminths, we coupled the single infection models for *T. retortaeformis* and *G. strigosum* through the immune response and examined different hypotheses of immune-mediated interaction that might occur between the two helminths. We assumed that helminth interactions may be mediated by antibody cross-immunity, where the effect of a specific antibody response produced against one helminth species also targets the other species (70). The full version of the dual infection model that accounts for all the helminth interactions and immune processes is the following:

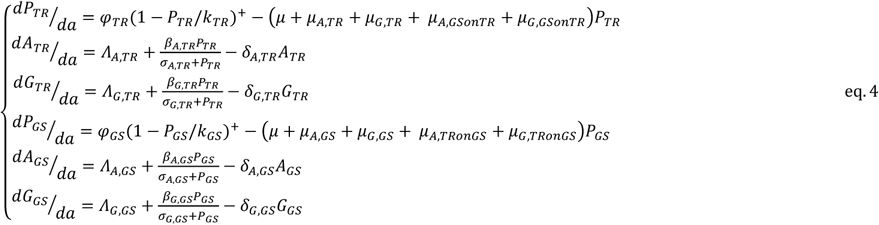

In Table 3, we reported the complete list and definition of model parameters. In addition to the components described for single infections (Table 1), *μ*_*A,GSonTR*_ and *μ*_*G,GSonTR*_ represent the cross-immunity attack rates of the specific IgA and IgG responses, respectively, stimulated by and produced against *G. strigosum* that also attack *T. retortaeformis*; vice versa, *μ*_*A,TRonGS*_ and *μ*_*G,TronGS*_ are the cross-immunity attack rates of *T. retortaeformis* antibodies that also attack *G. strigosum*. Here, the antibody attack rates are a function of the antibody levels (*A*_*TR*,_*A*_*GS*_, *G*_*TR*,_*G*_*GS*_) and their affinity to the specific helminth, i.e. Ig-specific attack rate (*μ*_*A,TR*,_*μ*_*G,TR*,_*μ*_*A,GS*_, *μ*_*G,GS*_), and the coinfecting helminth, i.e. Ig-cross attack rate (*μ*_*A,GSonTR*,_*μ*_*G,GSonTR*,_*μ*_*A,GSonTR*,_*μ*_*G,GSonTR*_). As in the case of Ig-specific attack rates, two hypotheses on antibody affinity were tested for the Ig-cross attack rates:

**Table 3.**
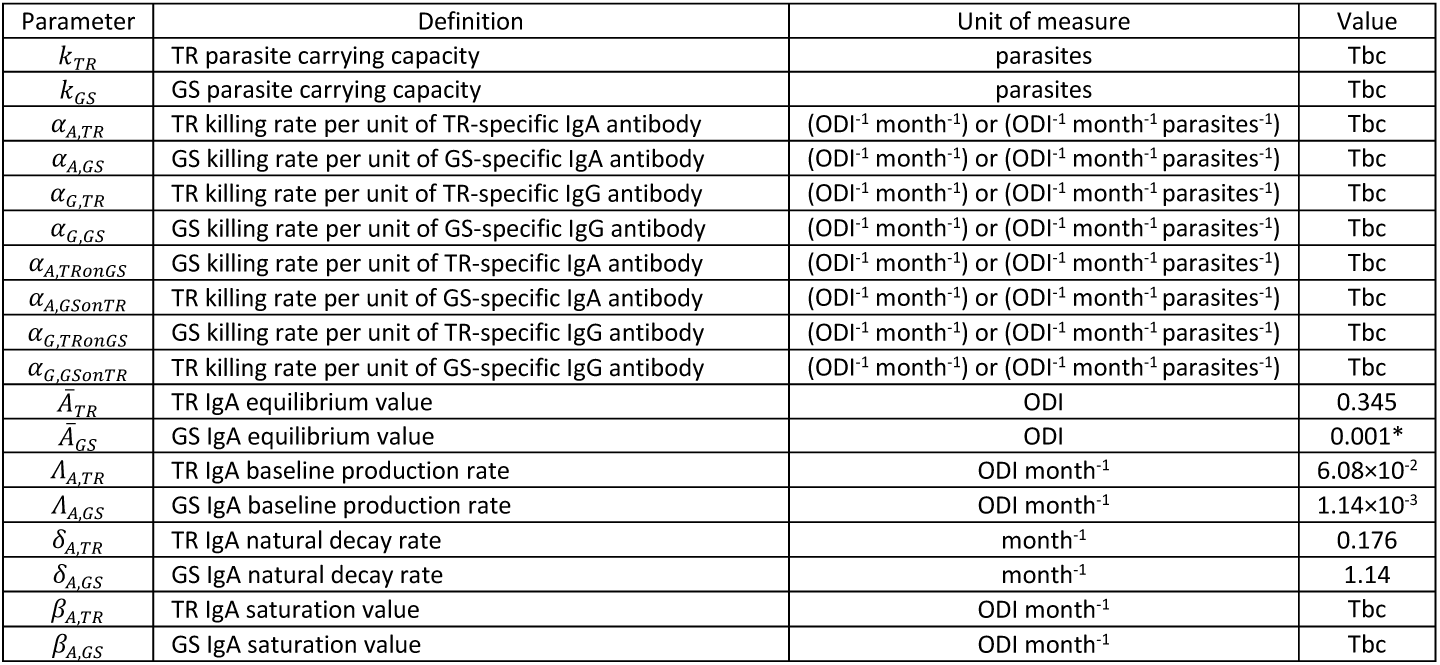
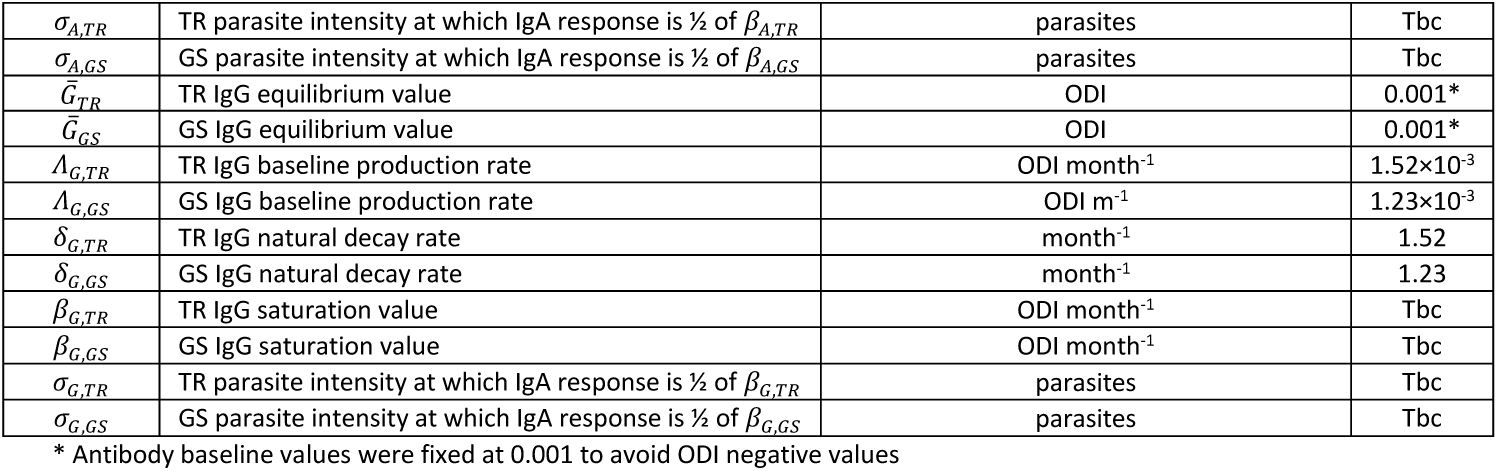
Dual infection: Model parameters, definitions, dimensions and available values for *T. retortaeformis* (TR) and *G. strigosum* (GS). Model parameters that are not reported in the table are assumed to take the same values as in the single infection case (see Table 1) (Tbc=to be calibrated).

i. a constant affinity by rabbit age to the cross-targeted species, *α*_*A,TRonGS*_, *α*_*G,TRonGS*_, *α*_*A,GSonTR*_, *α*_*G,GSonTR*,_ with the corresponding Ig-cross attack rates targeting parasite species *i* that are proportional to antibody levels produced against the other parasite species (*μ*_*A,TRonGS*_= *α*_*A,TRonGS*_*A*_*TR*_, *μ*_*A,GSonTR*_ = *α*_*A,GSonTR*_*A*_*GS*_, *μ*_*G,TRonGS*_= *α*_*G,TRonGS*_*G*_*TR*_, *μ*_*G,GSonTR*_ = *α*_*G,GSonTR*_*G*_*GS*_);
ii. an increasing affinity to the cross-targeted species by rabbit age, where the antibody affinity to the cross-target parasite species *i* is a function of the integral of exposure to the other helminth species from the birth of the rabbit up to its age *a* 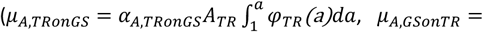 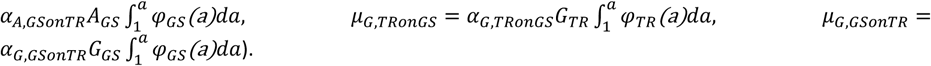. As previously noted for single infections, the baseline antibody production rates (*Λ*_*A,TR*,_*Λ*_*G,TR*,_*Λ*_*A,GS*_, *Λ*_*G,GS*_) and decay rates (*δ*_*A,TR*,_*δ*_*G,TR*_, *δ*_*A,GS*_, *δ*_*G,GS*_) were calculated from previous studies (57). To keep reasonable the dimensionality of the parameter-space to be explored, we assumed that the parameters describing the rates of rabbit feeding and exposure to infective parasite stages (*ϕ*_*TR*,_*ϕ*_*GS*_, *γ*_*TR*,_*γ*_*GS*_) were fixed to those identified as best models from single infections. While we could test different timing of infection between the two helminths, namely, *T. retortaeformis* followed by *G. strigosum* and vice versa, to reduce the complexity of our framework we assumed that the two helminths are ingested simultaneously by rabbits with both infections. This latter trend is probably the most common process occurring in natural conditions, as rabbits are constantly exposed to pasture contaminated with infective stages and carry persistent infections.

For the dual infection model, we recalibrated the parameters *α*_*A,TR*_, *α*_*G,TR*,_*k*_*TR*,_*α*_*A,GS*_, *α*_*G,GS*_ and *k*_*GS*_. These, together with the parameters that describe cross-immunity (*α*_*A,GSonTR*_, *α*_*G,GSonTR*_, *α*_*A,TRonGS*_, *α*_*G,TRonGS*_), were calibrated by minimizing the previously described loglikelihood function (see Appendix S1), simultaneously, for the two helminths, as:

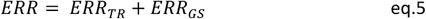

where

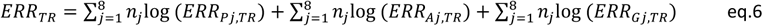

and

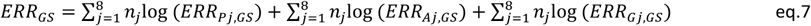

The error terms, *ERR*_*Pj,TR*_, *ERR*_*Aj,TR*_, *ERR*_*Gj,TR*_, *ERR*_*Pj,GS*_, *ERR*_*Aj,GS*_, and *ERR*_*Gj,GS*_, were computed as described for single infections (see eq.3 and ref. 26).

We evaluated our hypotheses of specific and cross-immunity, and the resulting seven competing models (Table 4), using *AIC* as discussed for single infections.

**Table 4.** Tested hypotheses and related mechanisms for the competing models of helminth interactions in dual infection. The parameters *α*_*A,TRonGS*_, *α*_*A,TRonGS*_, *α*_*G,TRonGS*_, *α*_*G,TRonGS*_ are set equal to 0 when the respective mechanism is not considered, otherwise they are calibrated (Tbc=to be calibrated). The additional hypotheses related to the species-specific components tested are listed in Table 2.

| Model | Hypotheses/Mechanisms | $\alpha_{Ai}$ | $\alpha_{Gi}$ |
| --- | --- | --- | --- |
| MD0 | Null model (no parasite immune-interaction) | 0 | 0 |
| MD1 | Age-dependent IgA-cross immunity (constant antibody affinity) | Tbc | 0 |
| MD2 | Age-dependent IgA-cross immunity (antibody affinity function of exposure) | Tbc | 0 |
| MD3 | Age-dependent IgG-cross immunity (constant antibody affinity) | 0 | Tbc |
| MD4 | Age-dependent IgG-cross immunity (antibody affinity function of exposure) | 0 | Tbc |
| MD5 | Age-dependent IgA-cross immunity + IgG-cross immunity (constant antibody affinity) | 0 | 0 |
| MD6 | Age-dependent IgA-cross immunity + IgG-cross immunity (antibody affinity function of exposure) | Tbc | Tbc |

### Confidence intervals of estimated parameters and sensitivity analysis

We assessed the 90% confidence intervals of the estimated parameters using a bootstrap procedure, reconstructing 100 replicates of the available time series by randomly sampling with replacement rabbit individual data of parasite intensities, IgA and IgG, and then calibrating for each replicate the parameter sets, minimizing the corresponding error function (71). We applied the described procedure to the models that minimized the *AIC* of the three infection groups *T. retortaeformis* single infection, *G. strigosum* single infection and dual infection. Additionally, to examine in more detail the relationship between infection and immunity in dual infections, we performed a sensitivity analysis of average parasite intensity across the eight age classes varying the parasite mortality rates induced by specific and cross-immunity (*e. g*. *α*_*A,GSonTR*_ and *α*_*A,TRonGS*_) from -75% to +75% of their calibrated values.

## Results

### Observed relationships in single and dual infections

We initially examined fundamental trends of the observed *T. retortaeformis* and *G. strigosum* intensities and antibody responses in rabbits with single and dual infections. The two helminths exhibited contrasting age-intensity relationships: *T. retortaeformis* intensities peaked in five months old rabbits and decreased as animals became older (Figure 1A), in contrast, *G. strigosum* intensities slowly increased with host age (Figure 1D). For both helminths, IgA and IgG levels increased with host age following a sigmoidal profile, reaching an upper asymptote in older hosts (Figure 1,B,C,E,F). A comparison between single- and dual-infected hosts (Table 5) showed that intensities were significantly (*p*<0.001) higher in dual-infected rabbits for both helminths (Figure 1A,D). Antibody levels were not significantly different between single and dual infections, however, when the interaction with age was taken into account IgG against *T. retortaeformis* was higher (*p*=0.029) whereas IgA against *G. strigosum* was lower (*p*=0.030) in rabbits with dual infections (Figure 1,B,C,E,F, Table 5).

**Table 5.** Generalized Linear Model (GLM) relating the observed parasite intensity, with a logarithmic link (negative binomial distribution), IgA and IgG levels (normal distribution) to host age (continuous variable), infection status (single/dual infection, categorical variable), and their interaction, for *T. retortaeformis* and *G. strigosum. n* represents the sample size.

| *T. retortaeformis*:<br>intensity | Coefficient | SE | p-value | *G. strigosum*:<br>intensity | Coefficient | SE | p-value |
| --- | --- | --- | --- | --- | --- | --- | --- |
| Intercept | 2.924 | 0.2067 | $< 2 \times 10^{-16}$ | Intercept | -5.055 | 0.3867 | $< 2 \times 10^{-16}$ |
| Single/Dual | 2.429 | 0.2883 | $< 2 \times 10^{-16}$ | Single/Dual | 3.221 | 0.4294 | $6.27 \times 10^{-14}$ |
| Age | 0.7630 | 0.0572 | $< 2 \times 10^{-16}$ | Age | 1.568 | 0.06411 | $< 2 \times 10^{-16}$ |
| Single/Dual*Age | -0.5048 | 0.06707 | $5.2 \times 10^{-14}$ | Single/Dual*Age | -0.5350 | 0.07165 | $8.2 \times 10^{-14}$ |
| *n* | 1386 |  |  | *N* | 1164 |  |  |
| *T. retortaeformis*:<br>IgA | Coefficient | SE | p-value | *G. strigosum*: IgA | Coefficient | SE | p-value |
| Intercept | 0.3461 | 0.1587 | 0.02932 | Intercept | -0.3514 | 0.2153 | 0.103 |
| Single/Dual | 0.003645 | 0.2214 | 0.9869 | Single/Dual | 0.4384 | 0.2758 | 0.112 |
| Age | 0.1404 | 0.04393 | 0.00143 | Age | 0.3404 | 0.04137 | $5.1 \times 10^{-16}$ |
| Single/Dual*Age | 0.02886 | 0.05151 | 0.5753 | Single/Dual*Age | -0.1111 | -2.173 | 0.030 |
| *n* | 1386 |  |  | *N* | 1164 |  |  |
| *T. retortaeformis*:<br>IgG | Coefficient | SE | p-value | *G. strigosum*: IgG | Coefficient | SE | p-value |
| Intercept | -0.1798 | 0.06456 | 0.00543 | Intercept | -0.1533 | 0.06921 | 0.0269 |
| Single/Dual | -0.08688 | 0.09009 | 0.33500 | Single/Dual | 0.02862 | 0.08867 | 0.7469 |
| Age | 0.1420 | 0.01787 | $4.1 \times 10^{-15}$ | Age | 0.1510 | 0.01330 | $< 2 \times 10^{-16}$ |
| Single/Dual*Age | 0.04582 | 0.02096 | 0.02897 | Single/Dual*Age | -0.006495 | 0.05112 | 0.6928 |
| *n* | 1386 |  |  | *n* | 1164 |  |  |

**Figure 1.**
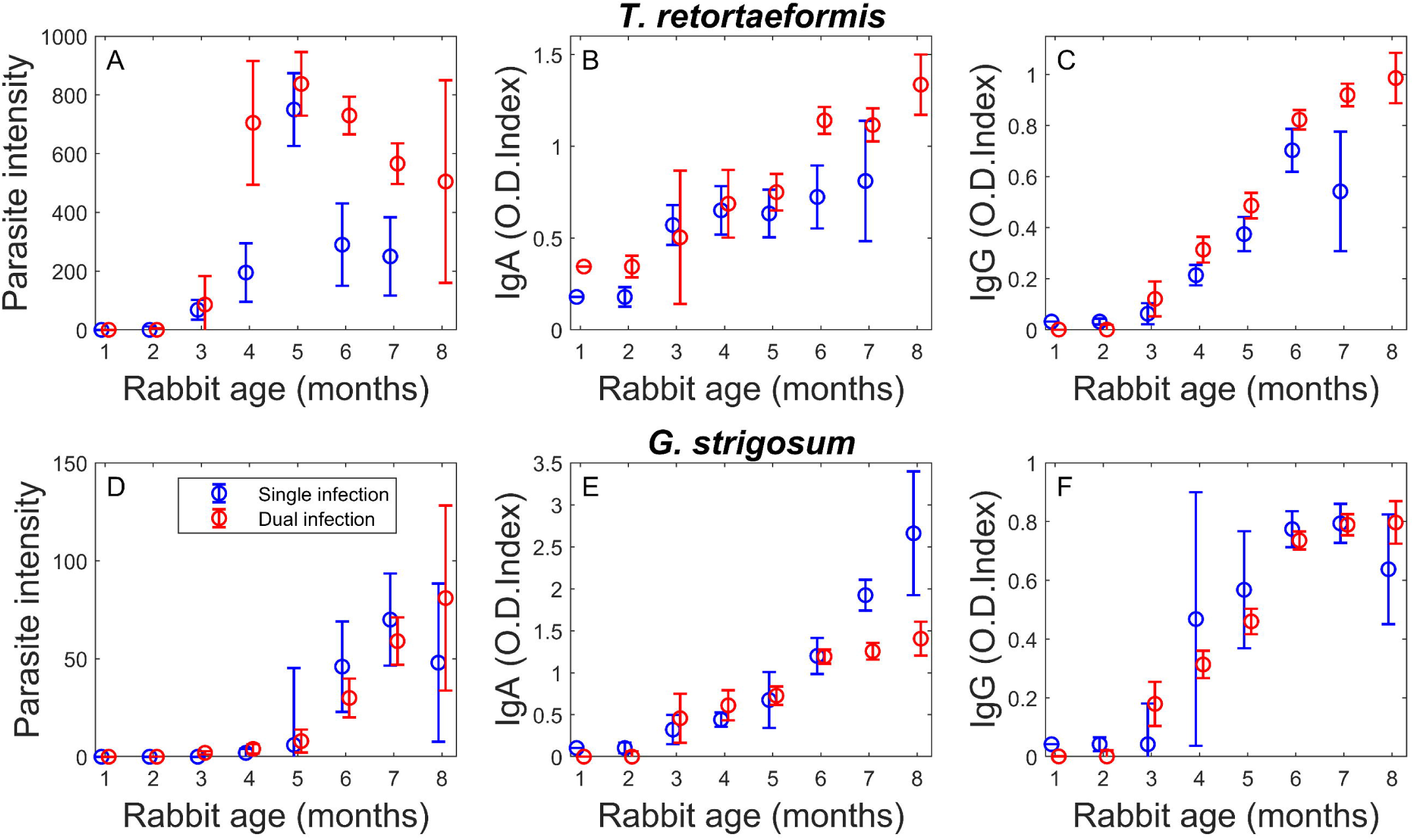
Observations of single (blue) and dual (red) infections. (A,D) Parasite intensity, (B,E) specie-specific IgA and (C,F) IgG response for *T. retortaeformis* (A-C) and *G. strigosum* (D-F) by rabbit age. Medians of observed data are represented with circles, while bars represent +/- 1 standard deviation ranges, estimated as +/- 1.4826 median absolute deviations.

### Model selection and simulation for single infections

Among the 14 independent models we tested, we found that parasite regulation by either IgA or IgG, with antibody affinity as a function of accumulated parasite exposure, could equally and well explain the dynamics of the two helminths in rabbits with single infections (models MS2 and MS4, *ΔAIC*<2, Table 6). When we considered the additive effect of both antibodies, model fitness did not increase (MS7, Table 6). This result suggests that while rabbits develop an antibody response which grows stronger with rabbit age due to increased antibody affinity, the impact of either IgA or IgG appears to be similar. For *G. strigosum*, the dynamics of infection could also be explained by parasite carrying capacity alone (MS5, Table 6), as model performance was comparable to the best models MS2 and MS4 (*ΔAIC*<2). However, if we base our evaluation strictly on model performance and select the model with the lowest *AIC* value, MS4 with IgG regulation best described *T. retortaeformis* single infection, while MS2 with IgA regulation best explained *G. strigosum* dynamics. The remaining competing models exhibited worse performances, although some *ΔAIC* were relatively close to the cut-off value of 2 but at the cost of a higher model complexity, as was the case of MS7 for both helminths. Here, we report the results of the selected models for each helminth in single infection (MS2 for *G. strigosum* and MS4 for *T. retortaeformis*), while we further analyzed the performance of the combined sub-optimal models (*ΔAIC*<2) when presenting the dual infections (see below and Table S1).

**Table 6.**
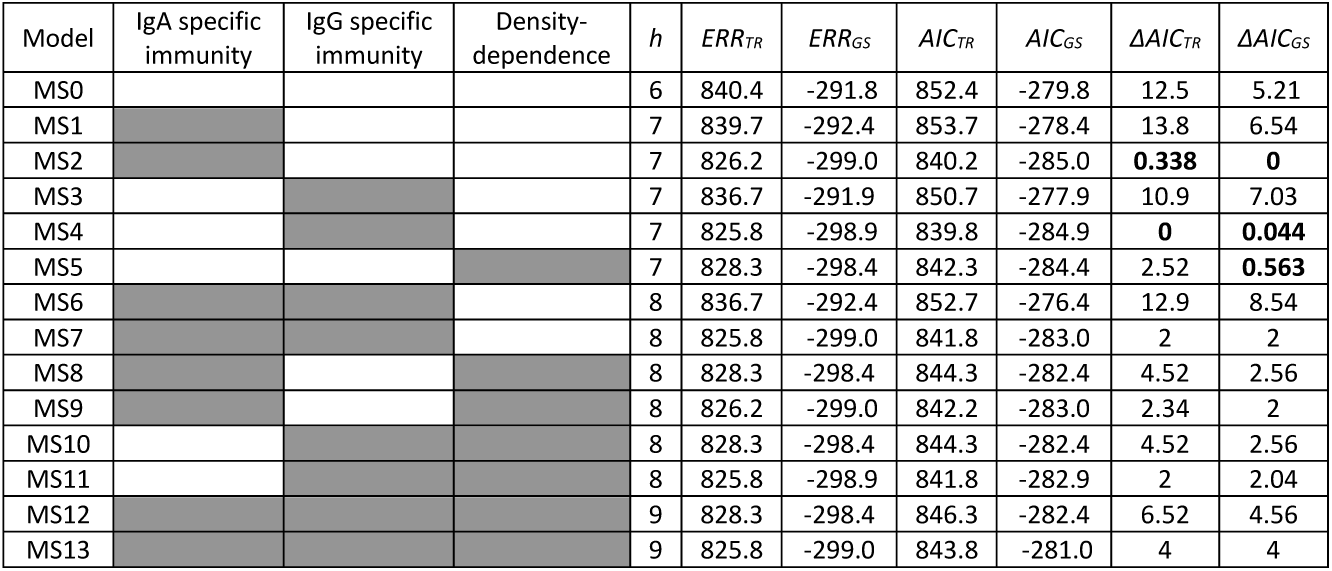
Summary of competing models for single infections based on level of complexity and performance. Tested models with activated regulation mechanisms (in grey), model complexity, *h*, error function, *ERR, AIC* and Δ*AIC scores* are reported for both *T. retortaeformis* (TR) and *G. strigosum* (GS). In bold are values of Δ*AICs* <2.

| Model | IgA specific immunity | IgG specific immunity | Density-dependence | $h$ | $ERR_{TR}$ | $ERR_{GS}$ | $AIC_{TR}$ | $AIC_{GS}$ | $\Delta AIC_{TR}$ | $\Delta AIC_{GS}$ |
| --- | --- | --- | --- | --- | --- | --- | --- | --- | --- | --- |
| MS0 |  |  |  | 6 | 840.4 | -291.8 | 852.4 | -279.8 | 12.5 | 5.21 |
| MS1 |  |  |  | 7 | 839.7 | -292.4 | 853.7 | -278.4 | 13.8 | 6.54 |
| MS2 |  |  |  | 7 | 826.2 | -299.0 | 840.2 | -285.0 | **0.338** | **0** |
| MS3 |  |  |  | 7 | 836.7 | -291.9 | 850.7 | -277.9 | 10.9 | 7.03 |
| MS4 |  |  |  | 7 | 825.8 | -298.9 | 839.8 | -284.9 | **0** | **0.044** |
| MS5 |  |  |  | 7 | 828.3 | -298.4 | 842.3 | -284.4 | 2.52 | **0.563** |
| MS6 |  |  |  | 8 | 836.7 | -292.4 | 852.7 | -276.4 | 12.9 | 8.54 |
| MS7 |  |  |  | 8 | 825.8 | -299.0 | 841.8 | -283.0 | 2 | 2 |
| MS8 |  |  |  | 8 | 828.3 | -298.4 | 844.3 | -282.4 | 4.52 | 2.56 |
| MS9 |  |  |  | 8 | 826.2 | -299.0 | 842.2 | -283.0 | 2.34 | 2 |
| MS10 |  |  |  | 8 | 828.3 | -298.4 | 844.3 | -282.4 | 4.52 | 2.56 |
| MS11 |  |  |  | 8 | 825.8 | -298.9 | 841.8 | -282.9 | 2 | 2.04 |
| MS12 |  |  |  | 9 | 828.3 | -298.4 | 846.3 | -282.4 | 6.52 | 4.56 |
| MS13 |  |  |  | 9 | 825.8 | -299.0 | 843.8 | -281.0 | 4 | 4 |

An analysis of the relationship between intensity of infection and antibody response highlighted further differences in the mechanisms of regulation affecting the contrasting dynamics of the two helminths observed in the field (Figures 1,2 and Table 7). Simulations of *T. retortaeformis* dynamics using the best model (MS4) suggested that IgG primarily controls the infection by reducing the burden in older rabbits while IgA is stimulated but does not affect parasite intensities. In fact, IgA stimulation by parasite intensity was turned on (*β*_*A,TR*_>0, *σ*_*A,TR*_ >0) but the impact on parasite intensity was turned off (*α*_*A,TR*_= 0; Figure 2A). Accordingly, both IgA and IgG responses were well described by sigmoidal functions of host age that reached their asymptotic values in older hosts (Figure 2B and C). The model underestimated the peak of parasite intensity, occurring in five-month-old rabbits, and the IgG peak at the host age of six months. In the first case it appears that the model was not flexible enough to describe this rapid increase. In the second situation the large IgG variation in seven-month-old rabbits, combined with a small sample size (N_7_=7), probably contributed to this trend. However, we want to stress that the model was fitted to individual data, while the simulated median-by-age values are presented in Figures 1 and 2.

**Table 7.** Single infections: Estimated values and 90% confidence intervals (CI) for the parameters of the selected models, MS4 for *T. retortaeformis* (TR) and MS2 for *G. strigosum* (GS). The 90% CIs are estimated via bootstrap.

| Parameter | TR value | GS value | TR 90%CI | GS 90%CI |
| --- | --- | --- | --- | --- |
| $\phi_i$ | $2.93 \times 10^4$ | $4.67 \times 10^3$ | $[9.16 \times 10^3; 1.47 \times 10^5]$ | $[4.46 \times 10^3; 1.33 \times 10^4]$ |
| $\gamma_i$ | 5.05 | 7.33 | $[4.17; 6.14]$ | $[7.32; 9.33]$ |
| $\alpha_{Ai}$ | - | $8.98 \times 10^{-3}$ | - | $[7.72 \times 10^{-3}; 1.54 \times 10^{-2}]$ |
| $\alpha_{Gi}$ | $5.30 \times 10^{-3}$ | - | $[2.66; 8.46] \times 10^{-3}$ | - |
| $\beta_{Ai}$ | 0.400 | 2.36 | $[0.328; 0.499]$ | $[2.21; 3.16]$ |
| $\sigma_{Ai}$ | 5.27 | 3.80 | $[2.29; 11.8]$ | $[1.43; 6.58]$ |
| $\beta_{Gi}$ | 1.10 | 0.584 | $[0.654; 3.66]$ | $[0.584; 0.691]$ |
| $\sigma_{Gi}$ | 177 | 0.239 | $[58.6; 1.03 \times 10^3]$ | $[3.44 \times 10^{-2}; 0.387]$ |

**Figure 2.**
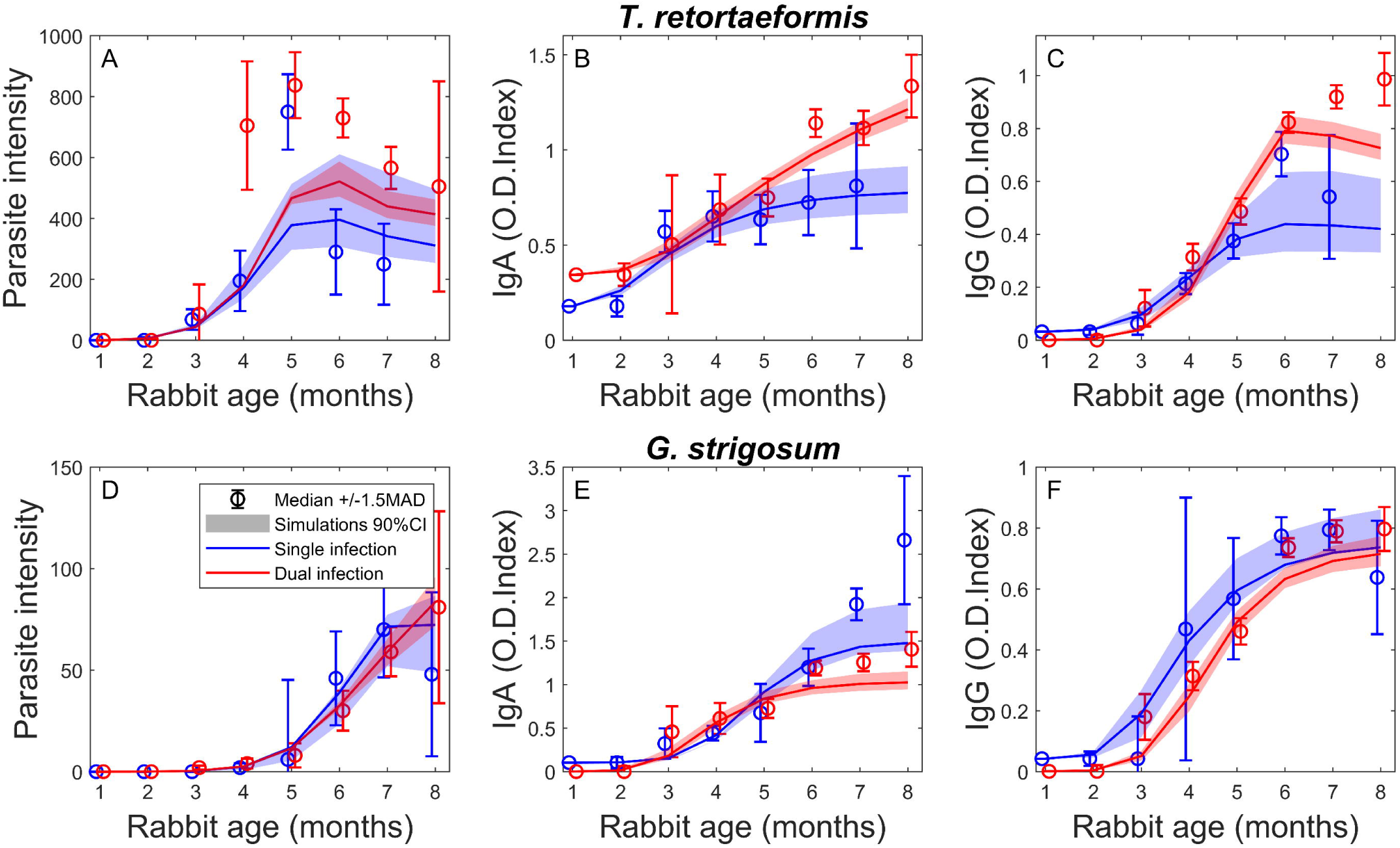
Observations and simulations of single (blue) and dual (red) infections. (A,D) Parasite intensity, (B,E) specie-specific IgA and (C,F) IgG response for *T. retortaeformis* (A-C) and *G. strigosum* (D-F) by rabbit age. Observed data (medians +/- 1.4826 median absolute deviations) are represented with circles and bars, while model simulations (MS4 for *T. retortaeformis* and MS2 for *G. strigosum* single infections, and MD4 for dual infections) with lines and shades, which correspond to the 90% confidence intervals.

The investigation of the stimulation and accumulation of antibodies against *T. retortaeformis* showed that while the IgA response was almost immediate and quickly reached the upper asymptote (*Λ*_*A,TR*_ + *β*_*A,TR*_) (Figure 3A, Table 3,7), the IgG response was much slower (*σ*_*A,TR*_ < *σ*_*G,TR*_) but increased with parasite intensity towards a higher asymptote (Figure 3C, Table 7). The ability of IgG to control *T. retortaeformis* was further examined by quantifying the IgG-specific parasite attack rate, calculated as the product of IgG level and affinity driven mortality by host age. We found that the attack rate was very low in rabbits up to the first four months of age but rapidly increased in older animals due to both higher antibody level but also stronger affinity and thus expected stronger mortality (Figure 4A).

**Figure 3.**
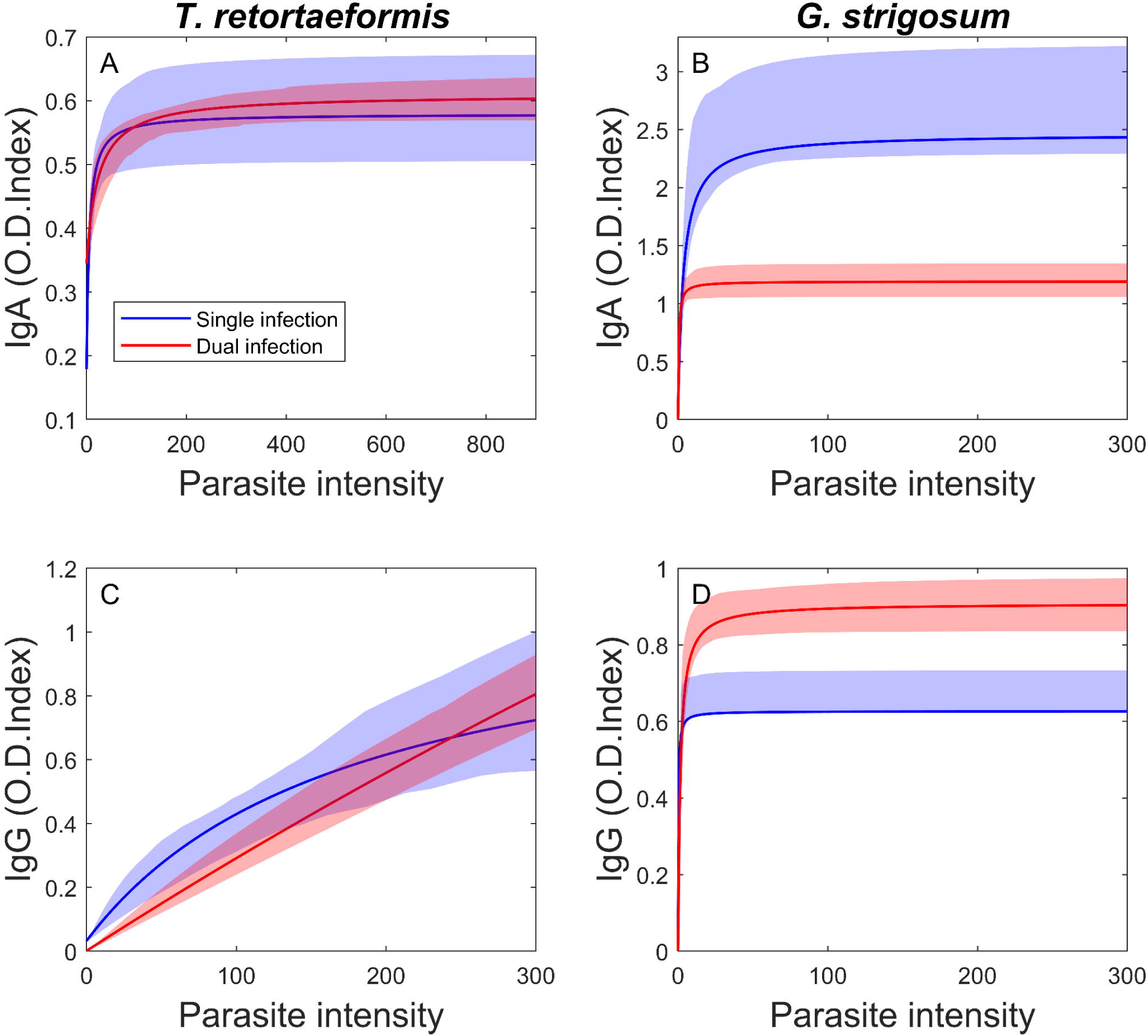
Antibody level (IgA: A,B; IgG: C,D) by parasite intensity for *T. retortaeformis* (A,C) and *G. strigosum* (B,D) in single (blue) and dual (red) infections. Lines with shades represent model simulations (MS4 for *T. retortaeformis* and MS2 for *G. strigosum* single infections; MD4 for dual infections) with the corresponding 90% CI evaluated via bootstrap.

**Figure 4.**
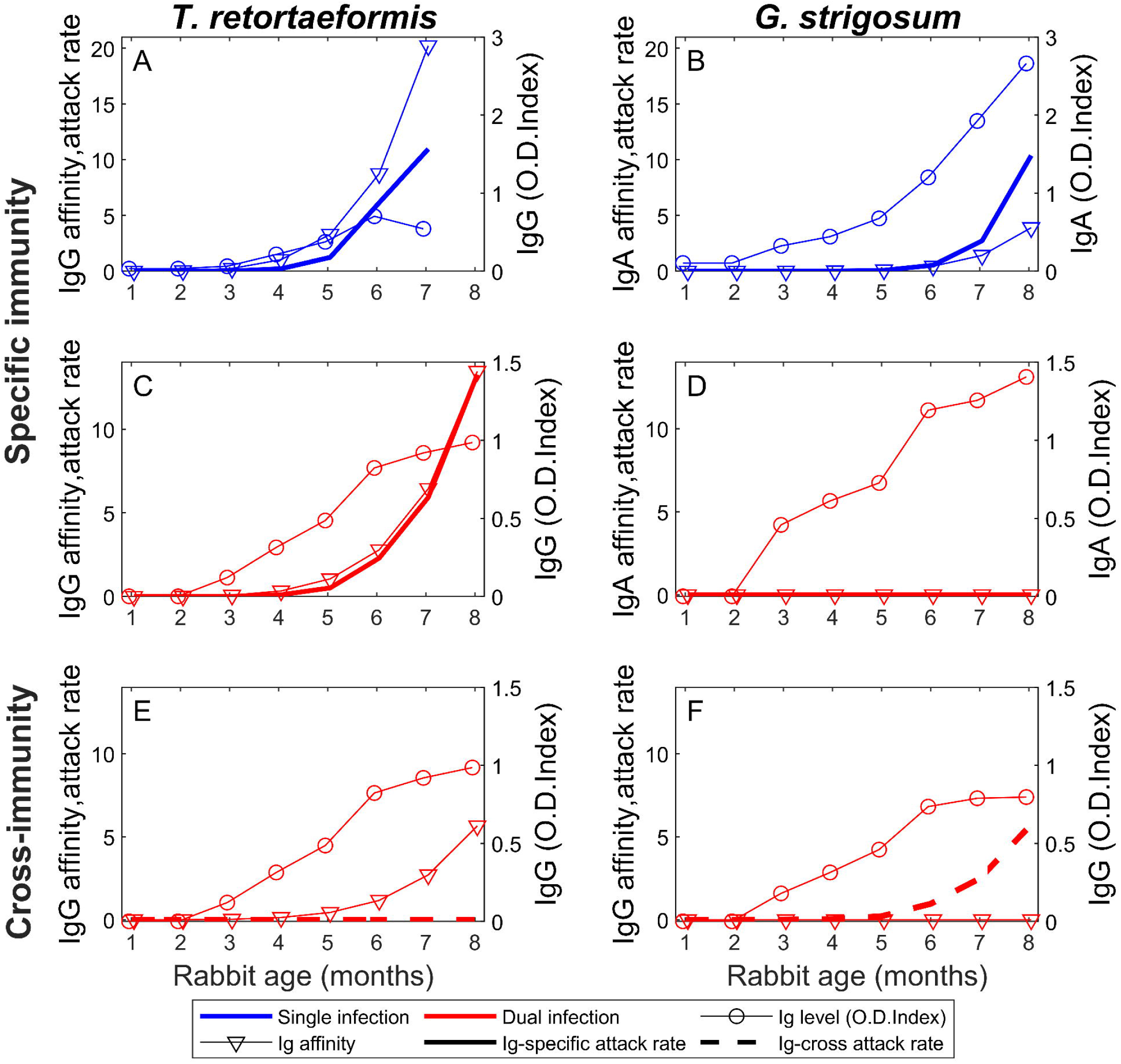
Antibody level, affinity and attack rate by specific (A,B,C,D) and cross- (E,F) immunity against *T. retortaeformis* (A,C,E) and *G. strigosum* (B,D,F by rabbit age, in single (A,B in blue) and dual (C,D,E,F in red) infections. For each rabbit age class, the median antibody level (circles, right axis), the median antibody affinity (triangles, left axis) and the median attack rate (bold lines, left axis) are reported. Ig attack rates are computed as the product between the median antibody level and affinity. Note that IgG-cross attack rate against *T. retortaeformis* (dashed line in panel E) is computed as the product of antibody level and affinity produced against *G. strigosum* (panel F) and vice versa for the IgG-cross attack rate against *G. strigosum* (dashed line in panel F).

For *G. strigosum* the best model MS2 selected IgA as the main antibody controlling the helminth, while IgG impact on parasite intensity was turned off (*α*_*G GS*_= 0) yet stimulated (*β*_*G,GS*_>0, *σ*_*G,GS*_>0). Simulations well reproduced the observed dynamics (Figure 2D-F), although there was a tendency to underestimate IgA in older age rabbits, probably because of the small sample size (*N*_*8*_=7). The investigation of the antibody stimulation and accumulation showed that both specific IgA and IgG responses to *G. strigosum* quickly saturated to the upper asymptote, with higher values for IgA than IgG (*Λ*_*A,GS*_+ *β*_*A,GS*_> *Λ*_*G,GS*_+ *β*_*G,GS*_) (Figure 3B,D and Tables 3,7) supporting the rapid antibody response of the rabbit to this chronic infection. However, the IgA-specific attack rate, which MS2 suggested to be responsible for the control of this infection, was very low in animals up to about six months of age but rapidly increased in 7/8-months old rabbits (Figure 4B). This indicates that the IgA response is probably not very effective during the first 5/6 months of age but could regulate *G. strigosum* abundance in older hosts that carried higher intensities (Figure 2D, Figure 4B). IgA levels in young rabbits were very low and the low IgA-specific attack rate in these age classes was probably caused by a low ability of antibodies to bind and kill parasites (Figure 4B). While the antibodies against the two helminths are not directly comparable because of their different characteristics, including their different dilutions, as a general trend we can speculate that the attack rate appeared to be faster to *T. retortaeformis* than *G. strigosum*.

### Model selection and simulation for dual infections

The models selected for single infections, MS4 for *T. retortaeformis* and MS2 for *G. strigosum*, were linked through a cross-antibody response and the type of cross-immunity that best described rabbits with dual infections was then examined by model selection. We identified MD4 as the best dual infection model, this included an expression of IgG cross-immunity in which species-specific IgG produced against one helminth also targeted the other helminth species and where the antibody affinity developed as a function of accumulated exposure to parasites (Table 8). From single infection models, MD4 included the species-specific IgG response against *T. retortaeformis* and the species-specific IgA response against *G. strigosum*. All the remaining models we tested had a *ΔAIC*>2, indicating that they were less suitable to parsimoniously describe the immune-mediated interactions between the two helminths. A possible exception to the above observation was model MD2, which included IgA cross-immunity and had a *ΔAIC* score very close to the cut-off value (*ΔAIC=*2.1).

**Table 8.** Summary of competing models for dual infection based on level of complexity and performance. Tested models with activated cross-regulation mechanisms (in grey), model complexity, *h*, error function for *T. retortaeformis, ERR*_*TR*_, and *G. strigosum, ERR*_*GS*_, total error function, *ERR, AIC* and Δ*AIC* scores are reported. In bold is value of Δ*AIC* <2.

| Model | IgA cross-immunity | IgG cross-immunity | $h$ | $ERR_{TR}$ | $ERR_{GS}$ | $ERR$ | AIC | $\Delta AIC$ |
| --- | --- | --- | --- | --- | --- | --- | --- | --- |
| MD0 |  |  | 10 | 1056 | 1931 | 2249 | 2269 | 2 |
| MD1 |  |  | 12 | 1056 | 1931 | 2249 | 2273 | 5.9 |
| MD2 |  |  | 12 | 1056 | 1189 | 2245 | 2269 | 2.1 |
| MD3 |  |  | 12 | 1056 | 1191 | 2247 | 2271 | 3.7 |
| MD4 |  |  | 12 | 1056 | 1187 | 2243 | 2267 | **0** |
| MD5 |  |  | 14 | 1056 | 1191 | 2247 | 2275 | 7.7 |
| MD6 |  |  | 14 | 1056 | 1187 | 2243 | 2271 | 4 |

In addition to the model selection just described and based on the best performing single infection models, we also examined alternative formulations for the species-specific antibody responses using models of single infection with ΔAIC<2 (Table 5), while varying the formulation of cross-immunity as well. Two structures were considered: *i*) IgA model, with IgA-specific immunity against each helminth, and *ii*) IgG model, with IgG-specific immunity against each helminth. For both settings, the best selected model of dual infection identified again IgG as cross-immunity (SI Table S1), confirming IgG as an important immune-mediating factor in the interactions between the two helminths, irrespective of the specific antibody isotype used.

The selected model, MD4, well captured the dynamics in dual-infected rabbits, particularly for *G. strigosum*; conversely, simulations tended to underestimate *T. retortaeformis* intensity and the related IgG levels in older age classes (Figure 2, Table 9), as already noted for single infections.

**Table 9.** Dual infection: Estimated values and 90% confidence intervals (CI) for the parameters of the selected model with IgG cross-immunity (MD4). The 90% CIs are estimated via bootstrap.

| Parameter | Value | 90%CI |
| --- | --- | --- |
| $\beta_{A,TR}$ | 0.256 | [0.231;0.300] |
| $\beta_{G,TR}$ | 5.48 | [3.08;8.48 x10<sup>6</sup>] |
| $\beta_{A,GS}$ | 1.19 | [1.09;1.35] |
| $\beta_{G,GS}$ | 0.907 | [0.855;0.991] |
| $\sigma_{A,TR}$ | 17.8 | [8.27;53.4] |
| $\sigma_{G,TR}$ | 1748 | [746;3.59x10<sup>9</sup>] |
| $\sigma_{A,GS}$ | 0.437 | [0.213;0.680] |
| $\sigma_{G,GS}$ | 1.52 | [1.05;2.38] |
| $\alpha_{A,GS}$ | 2.00x10<sup>-20</sup> | [0;3.78x10<sup>-4</sup>] |
| $\alpha_{G,TR}$ | 1.69x10<sup>-3</sup> | [1.69x10<sup>-3</sup>;2.61 x10<sup>-3</sup>] |
| $\alpha_{G,GSonTR}$ | 2.86x10<sup>-19</sup> | [0;5.51 x10<sup>-5</sup>] |
| $\alpha_{G,TRonGS}$ | 7.10x10<sup>-4</sup> | [6.16;9.74] x10<sup>-4</sup> |

*T. retortaeformis* intensity peaked in six-month-old hosts and decreased thereafter (Figure 2A), whereas IgA and IgG exhibited a sigmoidal response (Figure 2B,C). The investigation of IgA and IgG stimulation and accumulation by parasite intensity showed contrasting trends: while the IgA response reached its asymptotic value at low parasite intensities, the IgG response increase linearly but more slowly (Figure 3A,C). The IgG-specific attack rate against *T. retortaeformis* was low in young rabbits but markedly increased in adult hosts (Figure 4C), by contrast, the IgG-cross attack rate was fundamentally null due to low antibody affinity (Figure 4E). This result suggests that *T. retortaeformis* may be primarily regulated by a specific IgG response with minimal to no cross-effects by antibodies (Table 9). The sensitivity analysis of model MD4 allowed us to evaluate how *T. retortaeformis* intensities reacted to variations in the specific (*α*_*G,TR*_) and cross-immune (*α*_*G,GSonTR*_) responses (Figure 5A). Findings reinforced some of the results described above, namely by showing that specific IgG (*α*_*G,TR*_) appeared to be important for *T. retortaeformis* regulation, while IgG cross-immunity (*α*_*G,GSonTR*_) had a little effect. Finally, when we compared dual to single *T. retortaeformis* infections, simulations were able to capture the higher *T. retortaeformis* intensities observed, which could be explained by the combined effect of a low specific IgG attack rate (*α*_*G,TR DUAL*_ < *α*_*G,TR SINGLE*_, Tables 7,9, and Figure 4A,C) and a fundamentally ineffective cross-IgG reaction stimulated by *G. strigosum* but targeting *T. retortaeformis* (Figure 4E).

**Figure 5.**
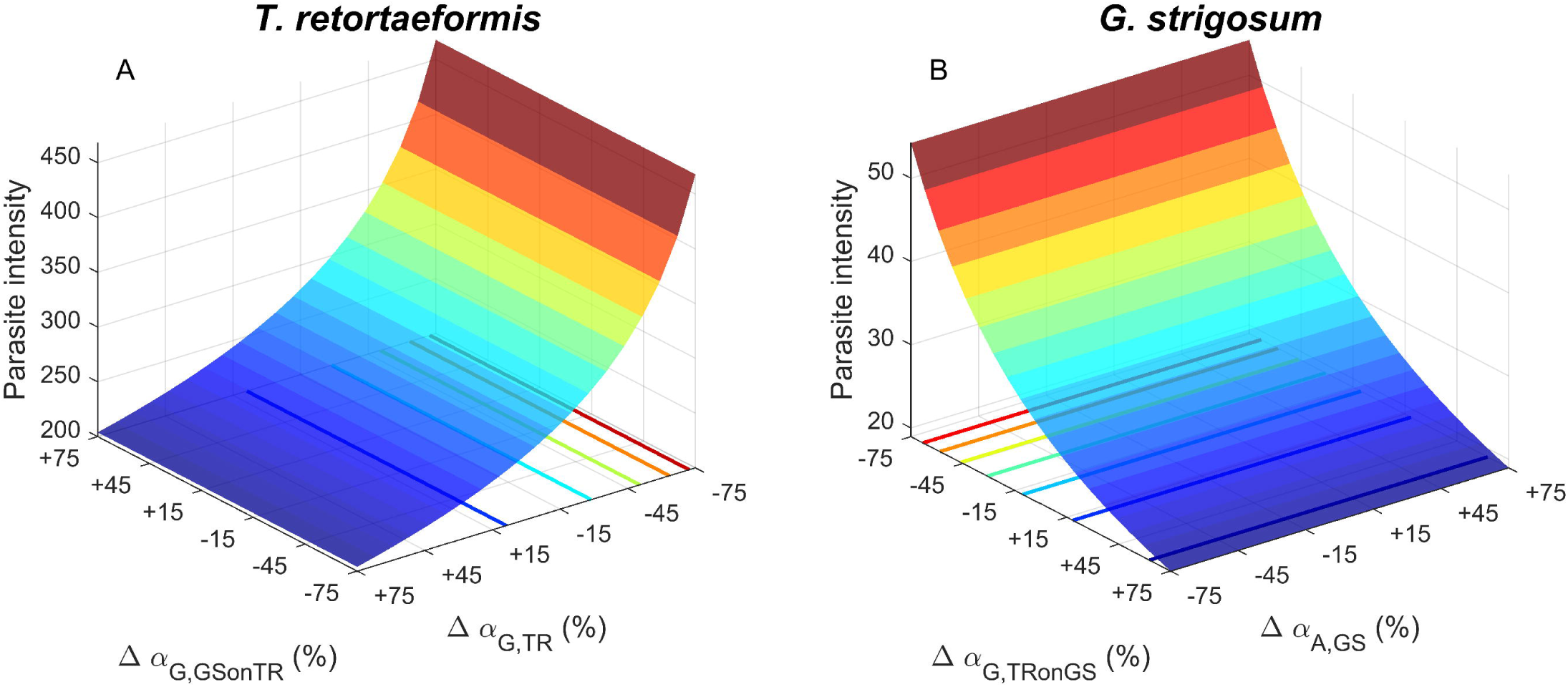
Sensitivity analysis of *T. retortaeformis* (A) and *G. strigosum* (B) average intensities by changes in parasite mortality rate induced by antibody specific (*α* _*G,TR*_ and *α*_*A,GS*_) or cross- (*α*_*G,GSonTR*_ and *α*_*G,TRonGS*_) responses. Simulations of parasite intensities were obtained from the selected model for dual infection MD4. Note that the axis directions differ in the two panels for better visualization and that contour lines use the same color code as the heat surfaces.

For *G. strigosum*, simulations well described parasite intensities (Figure 2D) and specific antibody responses, although there was a tendency to underestimate both antibody levels in older age classes (Figure 2E,F, Table 9). IgA and IgG stimulation by *G. strigosum* intensities was rapid, with different maximum values depending on the isotype (Figure 3B,D). Simulations suggested that the IgA-specific attack rate against *G. strigosum* in dual infection was negligible, which appeared not because of the low antibody production but because of their weak ability to bind and kill parasites (Figure 4D, Table 9). In contrast, IgG-cross attack rate stimulated by *T. retortaeformis* started to increase against *G. strigosum* at around six months of age (Figure 4F, Table 9). Model sensitivity to changes in the specific and cross-response against *G. strigosum* highlighted the negligible role of specific IgA (*α*_*A,GS*_) and showed that cross-IgG (*α*_*G,TRonGS*_) could cause significant changes in the intensities of this helminth for medium-to-high positive variation of the parameter values (Figure 5B). A visual comparison between single and dual infections found that the simulated *G. strigosum* intensities and antibody responses were lower in dual-compared to single-infected rabbits (Figure 2D-F) and reflected the empirical observation. In particular, the parasite-driven development of the IgA-specific response was slower and showed weak abilities to control the infection (*α*_*A,GSDUAL*_ < *α*_*A,GSSINGLE*_). Contrary to what observed for *T. retortaeformis*, this result suggests that *G. strigosum* could be primarily affected by an IgG cross-immune response, while the specific IgA response is essentially null. Ultimately, the asymmetric antibody reactions against the two helminth species could contribute to explain the contrasting dynamics of infection observed, as well as the mediating role of antibodies in their interactions from dual infected rabbits.

## Discussion

We developed a novel modeling framework that linked the response of IgA and IgG antibodies to the intensities of *T. retortaeformis* and *G. strigosum* in hosts infected either with one or two helminths to provide a quantitative understanding of the interactions between infection and host immunity in a natural population of rabbits. This study was motivated by the interest to improve the predicting capability of within-host models of single and dual infections by helminths and to explore if fundamental processes identified in a natural setting were consistent with studies of laboratory infections using the same host-helminth system.

We focused on antibodies, an immune variable that should be easy to measure in the field and which relative contribution of specific and cross-reacting responses can be estimated to quantify host-parasite and parasite-parasite interactions. While confirming general knowledge on their regulatory role, our mathematical framework provides a way forward to the measurement of first, antibody attack rate and its relationship with antibody affinity and second, the relative contribution of antibody specific and cross-responses in hosts with co-infections. Whilst these measurements need to be corroborated with molecular analysis our findings offer a biologically meaningful and testable explanation. Model selection suggested that, although IgG is selected against *T. retortaeformis* and IgA against *G. strigosum*, both antibodies could well explain the dynamics of the two helminths in rabbits with single infections. For rabbits with dual infections, we found that the two helminths interacted via asymmetric IgG cross-immunity, which was stronger against *G. strigosum* than *T. retortaeformis*, irrespective of the antibody isotype specific to each species. Overall, we showed that differences in antibody affinity and attack rate could explain the contrasting dynamics of the two helminths in both single and dual infections.

In several host-parasite systems, like *Ostertagia circumncincta* and *Ostertagia leptospicularis* in sheep, *Fasciola hepatica* and *Cooperia oncophora* in cattle and *Trichinella britovi* in mice, the species-specific responses of IgA, IgG, and IgE have frequently been associated with the regulation of parasite intensities (72–80). However, whether and which of these isotypes were actively controlling parasite dynamics was unclear. Our modelling approach represents a useful tool to disentangle the relative contribution of these antibodies, including their cross-reacting role in hosts with multiple infections.

We found that model selection consistently recommended the framework where the attack rate of antibodies was defined as a function of the accumulated host exposure to past and current infections of the target helminth. This finding is in agreement with previous models that conceptualized acquired immunity as a functional response to the history of infection (16,18,20,22,23,27). We further examined the antibody response and found that its strength was determined by the level of antibodies in blood serum as well as by their ability to kill parasites, which we defined as antibody affinity. By matching empirical data on antibodies and infection, our simulations were able to provide a more accurate description of how the relationship between these two variables could change with host age, and thus how the accumulated exposure to an infection could affect antibody affinity over time. This also allowed us to quantify the non-linear properties of species-specific antibody attack rates, which were extremely low in newborns and juveniles but quickly increased in adults as a response to the cumulated exposure to infective stages of both helminths in single infections. This rapid increase of the antibody attack rate in adults could explain the decrease of *T. retortaeformis* intensities in older rabbits, albeit it did not confer long-term protection to reinfection. In contrast, the accumulation of *G. strigosum* in adults was probably caused by a slow increase of antibody affinity, which was only apparent in older hosts, although they were still incapable to effectively control the helminth.

The relationship between parasite intensity and parasite killing by the immune response could also be represented by the typical functional responses used to describe a predator-prey relationship in ecology (40). Our empirical data indicated that both IgA and IgG exhibited a sigmoidal profile by host age and our model assumption was that the antibody response (predator) developed as a saturating function of parasite load (prey) (81,82). Field data confirmed these saturating antibody-infection trajectories, except for IgG to *T. retortaeformis*; these trajectories also showed differences in their strength both between helminth species and between single and dual infections. We suggest that these differences, and the resulting dynamics of infection of the two helminths, could be explained by contrasting attack rates by species-specific antibodies, as noted above.

When *T. retortaeformis* and *G. strigosum* coinfect the same host, simulations well described the relative changes of helminth intensity and related antibodies observed in the field and provided a conceivable mechanism for the mediating role played by IgA and IgG in helminth interactions. The higher intensities observed in dual than single infections could be explained by a lower specific IgG attack rate to *T. retortaeformis* and a specific IgA attack rate to *G. strigosum* that was fundamentally null. Higher intensities were also affected by an asymmetry in helminth immune-mediated interactions generated by differences in antibody cross-attack rate and related cross-affinity, i.e. the ability of recognizing the second helminth species by antibodies produced against the first species. Specifically, antibody cross-affinity was negligible against *T. retortaeformis* but more apparent against *G. strigosum* in old rabbits. Therefore, the combination of a null specific IgA attack rate and a moderate IgG cross-attack rate against *G. strigosum* were arguably insufficient to control this helminth resulting in a constant accumulation in older hosts. For *T. retortaeformis*, a low specific IgG attack rate together with a negligible IgG cross-attack rate from *G. strigosum* antibodies could explain the high intensities and most of the dynamics in dual infected rabbits.

The general findings from the current study were consistent with previous modeling work using laboratory data, where additional complexities in the immune response were examined by including the intermediate role of interleukin 4 (IL4) to IgA production (48). Specifically, the first common result was that differences in the strength of the immune response, rather than changes of the immune mechanism, could explain the contrasting dynamics of the two helminths, irrespective of the type of infection. The second common finding was the occurrence of an asymmetric antibody cross-response that was mainly directed against *G. strigosum* and null against *T. retortaeformis* in rabbits with both helminths. In this study the focus was on IgA cross-immunity, however, we currently showed that although IgG cross-immunity was consistently selected in our models the alternative framework with IgA was very similar (MD2: ΔAIC=2.1), suggesting that the two antibodies might have a similar role. Indirect interactions via antibody cross-immunity have been proposed to explain parasite coinfections in other animal systems from natural and experimental settings (83,84). For example, IgA, IgG, and IgE have been shown to be involved in helminth interactions and their response has been found to change in presence of a second helminth species (61,85). Our work provided a quantitative explanation of how these interactions could occur in the rabbit-helminth system and likely in other host-parasite settings.

We have proposed a quantitative, yet tractable modeling framework to describe the fundamental characteristics of the antibody response to, and the infection dynamics of, two gastrointestinal helminths. A novel aspect of our model is the estimation of antibody attack rate and affinity both in single and dual infections, beside the general description of the antibodies produced against each helminth. The quantification of antibody affinity maturation is crucial to better understand the immune-infection relationship and its changes throughout the host lifespan. Repeated exposure to the same antigen stimulates the production of antibodies with a greater affinity as a result of clonal selection occurring at the germinal center of B cells and somatic hypermutations (86). We used mathematical modeling to quantify antibody affinity maturation by host age, for both specific and cross-immunity in single and dual infections, however, further laboratory analyses are needed to confirm our findings and identify within-host interactions that could influence the development of this affinity.

The immune response to helminths is complex (78,87–89) and other immunological components and relations, in addition to those considered in this study, are expected to contribute to the observed patterns (48,90). We focused on antibodies that are important and well characterized in parasite immunology and provided a plausible explanation of how IgA and IgG interact with two helminth infections. Our within-host model successfully described the dynamics observed in the field but omitted to consider temporal changes in the availability of free-living infective stages on the pasture, which is required to quantify seasonal variation in host exposure. Therefore, we cannot exclude that some of the variations observed in the intensity of infection at specific age classes was affected by temporal changes in the exposure of rabbits to the pool of free-living stages. This issue and the role of climatic drivers in the availability of free-living stages is currently being investigated (46).

Our study provides novel insights into a quantitative understanding of immune-infection relationships in natural hosts with single and dual helminth infections. By explicitly incorporating measurements of the antibody responses and infection by host age, we were able to disentangle some of the heterogeneities that characterize our rabbit-helminth system and that are expected to be common in many other natural animal settings. The complexity and reliability of these models ultimately depends on the availability of empirical data. There is a long history of modeling the dynamics of helminth infections, the next critical step is to develop mechanistic models that can accurately capture some of the within-host complexities in the interactions between parasites and host immunity and embrace the evidence that these processes often occur in hosts infected by more than one parasite species.

## Supporting information

**Appendix S1. Akaike criterion with lognormally distributed noise**

**Table S1. Summary of competing models for dual infections based on performance and level of complexity**. Tested models with activated cross-regulation mechanisms (in grey), model complexity, *h*, error function, *ERR*, and *AIC* and Δ*AIC* scores. Candidate models were built by combining the suboptimal model structures for each helminth in the single infection model selection, i.e. MS2 and MS4.

## Acknowledgments

This study was supported by the National Science Foundation (DEB-0716885 and DEB-1145697).

## Author contributions

**Conceptualization:** Isabella M. Cattadori, Chiara Vanalli, Lorenzo Mari, Renato Casagrandi, Marino Gatto.

**Data curation:** Isabella M. Cattadori.

**Formal analysis:** Chiara Vanalli, Lorenzo Mari.

**Funding acquisition:** Isabella M. Cattadori.

**Investigation:** Chiara Vanalli, Renato Casagrandi, Marino Gatto, Isabella M. Cattadori.

**Methodology:** Lorenzo Mari, Marino Gatto, Renato Casagrandi, Chiara Vanalli.

**Project administration:** Isabella M. Cattadori.

**Supervision:** Isabella M. Cattadori.

**Validation:** Chiara Vanalli.

**Visualization:** Chiara Vanalli.

**Writing – original draft:** Chiara Vanalli, Isabella M. Cattadori.

**Writing – review & editing:** Chiara Vanalli, Lorenzo Mari, Renato Casagrandi, Marino Gatto, Isabella M. Cattadori.

