## Supplementary material for "Modeling the contribution of antibodies to the within-host dynamics of single and dual helminth infections in a natural system": S1 Appendix

**Appendix S1. Akaike criterion with lognormally distributed noise**

Assume data $X_i$, where $i$ = 1, …$n$ is time or age, are affected by uncorrelated lognormal noise $\varepsilon_i$, i.e. $X_i = \tilde{X}_i \varepsilon_i$ with ln($\varepsilon_i$) being normally distributed with 0 mean and unknown variance $\sigma^2$. Let $\tilde{X}_i$ be a function of $i$ and be defined by a vector $\theta$ of $h$ unknown parameters. Therefore

$$\ln\left(X_i/f(i;\theta)\right) = \ln\left(\varepsilon_i\right)$$

and the likelihood function is

$$L(X;\theta,\sigma) = \prod_{i=1}^{n}\left[(2\pi\sigma^2)^{-\frac{1}{2}}X_i^{-1}exp\left(-\frac{(\ln\left(X_i/f(i;\theta)\right))^2}{2\sigma^2}\right)\right]$$

$$=(2\pi\sigma^2)^{-\frac{n}{2}}exp\left(-\frac{\sum_{i=1}^{n}(\ln\left(X_i/f(i;\theta)\right))^2}{2\sigma^2}\right)\prod_{i=1}^{n}X_i^{-1}$$

The loglikelihood is thus

$$\mathcal{L}(X;\theta,\sigma) = -\frac{n}{2}\ln(2\pi\sigma^2) - \sum_{i=1}^{n}\ln(X_i) - \frac{\sum_{i=1}^{n}(\ln\left(X_i/f(i;\theta)\right))^2}{2\sigma^2}$$

and is maximized by finding the estimated value $\hat{\sigma}^2$ of the variance and $\hat{\theta}$ of $\theta$ that maximize the loglikelihood. For any given $\hat{\theta}$ the optimal value of the variance $\sigma^2$ is provided by the formula

$$\hat{\sigma}^2 = \frac{\sum_{i=1}^{n}\left(\ln\left(X_i/f\left(i;\hat{\theta}\right)\right)\right)^2}{n}$$

The resulting loglikelihood is thus

$$\mathcal{L}\left(X;\hat{\theta},\hat{\sigma}\right) = -\frac{n}{2}\ln(2\pi) - \frac{n}{2}\ln(\hat{\sigma}^2) - \sum_{i=1}^{n}\ln(X_i) - \frac{n\hat{\sigma}^2}{2\hat{\sigma}^2} = const - \frac{n}{2}\ln(\hat{\sigma}^2)$$

where $const = -\frac{n}{2}\ln(2\pi) - \sum_{i=1}^{n}\ln(X_i) - \frac{n}{2}$. Therefore the optimal $\hat{\theta}$ is the one that minimizes $\hat{\sigma}^2$ namely the squared error $\sum_{i=1}^{n}(\ln\left(X_i/f(i;\theta)\right))^2$. By discarding the constant and multiplying by 2 one obtains the Akaike criterion index, namely

$$AIC = n\ln\left(\frac{\sum_{i=1}^{n}\left(\ln\left(X_i/f\left(i;\hat{\theta}\right)\right)\right)^2}{n}\right) + 2h$$

Whenever we have data $X$, $Y$, $Z$ etc. each with its own variance and number of data, under the assumption of independence of the noises affecting each set of data, AIC generalizes (easy to prove following the same way of reasoning illustrated above) to

$$AIC = n_X\ln\left(\frac{\sum_{i=1}^{n}\left(\ln\left(X_i/f_X\left(i;\hat{\theta}\right)\right)\right)^2}{n_X}\right) + n_Y\ln\left(\frac{\sum_{i=1}^{n}\left(\ln\left(Y_i/f_Y\left(i;\hat{\theta}\right)\right)\right)^2}{n_Y}\right)$$

$$+n_Z\ln\left(\frac{\sum_{i=1}^{n}\left(\ln\left(Z_i/f_Z\left(i;\hat{\theta}\right)\right)\right)^2}{n_Z}\right) + 2h$$
