## Supplementary material for "Modeling the contribution of antibodies to the within-host dynamics of single and dual helminth infections in a natural system": S1 Table

**Table S1. Summary of competing models for dual infections based on performance and level of complexity.** Tested models with activated cross-regulation mechanisms (in grey), model complexity, *h*, error function, *ERR*, and *AIC* and Δ*AIC* scores. Candidate models were built by combining the suboptimal model structures for each helminth in the single infection model selection, i.e. MS2 and MS4.

<table>
<tr><th>Single infection T. retortaeformis</th><th>Single infection G. strigosum</th><th>Tested model</th><th>IgA cross-immunity</th><th>IgG cross-immunity</th><th>h</th><th>ERR</th><th>AIC</th><th>ΔAIC</th></tr>
<tr><td rowspan="7">MS2: IgA specific-immunity</td><td rowspan="7">MS2: IgA specific-immunity</td><td>MD0</td><td></td><td></td><td>10</td><td>2251.9</td><td>2271.9</td><td>2.6</td></tr>
<tr><td>MD1</td><td style="background:grey"></td><td></td><td>12</td><td>2251.9</td><td>2275.9</td><td>6.6</td></tr>
<tr><td>MD2</td><td style="background:grey"></td><td></td><td>12</td><td>2248.2</td><td>2272.2</td><td>2.9</td></tr>
<tr><td>MD3</td><td></td><td style="background:grey"></td><td>12</td><td>2248.1</td><td>2272.1</td><td>2.8</td></tr>
<tr><td>MD4</td><td></td><td style="background:grey"></td><td>12</td><td>2245.3</td><td>2269.3</td><td>0</td></tr>
<tr><td>MD5</td><td style="background:grey"></td><td style="background:grey"></td><td>14</td><td>2248.1</td><td>2276.1</td><td>6.8</td></tr>
<tr><td>MD6</td><td style="background:grey"></td><td style="background:grey"></td><td>14</td><td>2245.3</td><td>2273.3</td><td>4</td></tr>
<tr><td rowspan="7">MS4: IgG specific-immunity</td><td rowspan="7">MS4: IgG specific-immunity</td><td>MD0</td><td></td><td></td><td>10</td><td>2249.5</td><td>2269.5</td><td>3.4</td></tr>
<tr><td>MD1</td><td style="background:grey"></td><td></td><td>12</td><td>2249.0</td><td>2273.0</td><td>6.9</td></tr>
<tr><td>MD2</td><td style="background:grey"></td><td></td><td>12</td><td>2244.3</td><td>2268.3</td><td>2.2</td></tr>
<tr><td>MD3</td><td></td><td style="background:grey"></td><td>12</td><td>2245.8</td><td>2269.8</td><td>3.7</td></tr>
<tr><td>MD4</td><td></td><td style="background:grey"></td><td>12</td><td>2242.1</td><td>2266.1</td><td>0</td></tr>
<tr><td>MD5</td><td style="background:grey"></td><td style="background:grey"></td><td>14</td><td>2245.8</td><td>2273.8</td><td>7.7</td></tr>
<tr><td>MD6</td><td style="background:grey"></td><td style="background:grey"></td><td>14</td><td>2242.1</td><td>2270.1</td><td>4</td></tr>
</table>
